# DivQuant: Estimation of Species Richness and Entropy from Small Samples

**DOI:** 10.64898/2026.06.08.730836

**Authors:** Johanna Elena Schmitz, Sven Rahmann

## Abstract

Estimating diversity properties of discrete distributions from a small observed sample is a fundamental problem in algorithmic statistics that has applications in many fields, in particular bioinformatics, but also in ecology or linguistics. The two most common diversity measures are the number of distinct elements in a multiset, also referred to as “species richness” in ecology or “alpha diversity” in microbial analysis, and the Shannon entropy, also referred to as “evenness”. Estimating these properties from a small sample is particularly challenging for distributions with many rare elements. Thus, many estimators have been proposed in the past that, in practice, work well for different types of distributions.

We present DivQuant, an optimization-based, extrapolating richness and entropy estimator with three contributions. First, we formulate the upsampling problem as a convex quadratic program with a Neyman *χ*^2^ objective. Unlike the linear program of its predecessor RichnEst, DivQuant admits confidence intervals via *χ*^2^ test inversion that are empirically well-calibrated. Second, we replace RichnEst’s fixed-threshold fingerprint truncation with the rare/abundant fingerprint split of Valiant and Valiant, which strongly reduces problem size and preserves enough degrees of freedom for the confidence-interval program to remain valid and feasible. Third, we plug the optimal population fingerprint returned by the program into Shannon’s entropy formula to obtain an entropy estimate. DivQuant attains close-to-nominal 95% confidence intervals in essentially all tested regimes, including six simulated distribution families, *Tara Oceans* microbiome data, and *10X Genomics* scRNA-seq data, while competing state-of-the-art methods (RichnEst, iNext, PreSeq) miss the true richness in up to 80% of instances, well above the nominal 5%. In addition, DivQuant outperforms classical asymptotic entropy estimators (Miller-Madow, CAE) and the extrapolating iNext estimator.

Running times remain competitive, with DivQuant typically completing in seconds. DivQuant is available as a command-line tool at https://gitlab.com/rahmannlab/divquant.

**2012 ACM Subject Classification:** Mathematics of computing *→* Probability and statistics; Mathematics of computing *→* Linear programming; Mathematics of computing *→* Quadratic programming; Applied computing *→* Bioinformatics

## 1 Introduction

With access to only a small sample from a large population, there are many questions one may ask about the total population. For example, how many distinct elements went unseen in the sample? Or, how even is the population, i.e., what is the composition of the population regarding rare and abundant elements? The term *richness* is often used to denote the number of distinct elements in a population or sample. Evenness is measured using *entropy*, often referred to as Shannon diversity [38]. Both measures are important diversity measures in many fields, such as ecology [16, 41], linguistics [19, 20] and biology [5]. In microbial sequence analysis, richness (also called alpha diversity) and entropy are commonly used to analyze bacterial communities or environments [43, 40], where both function and community dynamics are assessed based on diversity changes over time. Furthermore, both measures are used to evaluate the diversity of immune repertoires or to track changes in the immune system during health and disease [18, 34].

However, there is an open debate about how best to compare the diversity between samples and which diversity measure to use [6]. Richness is usually computed using either the observed sample richness, which underestimates the true richness for almost all samples, or by using a statistical estimate. The same is true for entropy. An overview of the sampling problem is shown in Figure 1A–C.

**Figure 1.**
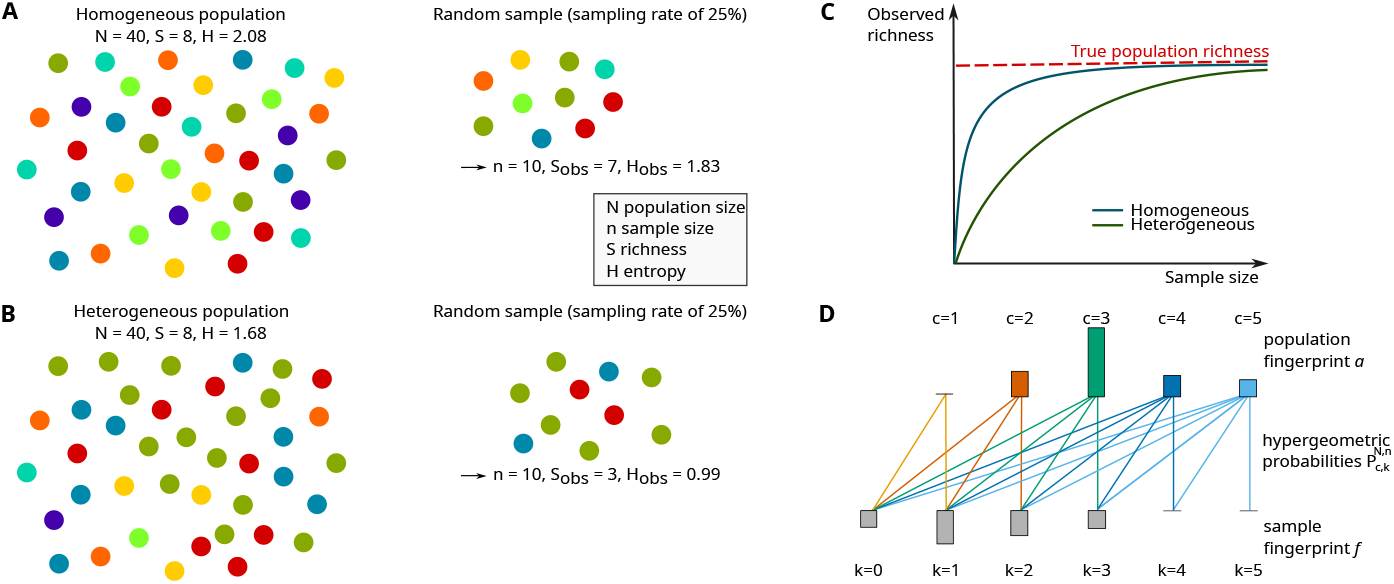
Difference between population richness *S* (entropy *H*) and observed sample richness (entropy). Colors denote distinct species. **A** In a homogeneous population, a small sample suffices to observe a richness close to the true value. **B** For a population with many rare species, the observed sample richness underestimates the population richness. **C** Species richness is commonly assessed via a rarefaction curve: as the sample size grows, the observed richness converges to the population richness, at a rate that depends on the species composition. **D** Example of the downsampling process from a population fingerprint *a* to a sample fingerprint *f*. A species observed 3 times in the sample must have abundance *≥* 3 in the population, so the expected value of *f*_3_ is *a*_3_ *·*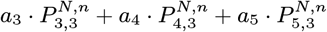 (see Section 2.2). Figure adapted from [36].

### Related work

Richness and entropy estimators can be divided into two groups depending on whether they require the full population size to be known or not, referred to as extrapolating and asymptotic estimators, respectively [9]. Ecological studies typically require accurate asymptotic lower bounds or estimates for extrapolated populations that are up to three times larger than the observed sample. Other applications, such as metagenomics or immune repertoire diversity, require accurate estimates for populations that are more than ten times larger than the observed sample.

Some asymptotic estimators, like the Chao 1 [8] or Jackknife estimator [7], assume that the number of species seen only once or twice is most predictive of the number of unseen species. Other estimators assume that the sample was drawn from a parametric family of probability distributions, e.g., a Gamma-Poisson mixture distribution, and reduce the problem to estimating the parameters of a parametric probability distribution, using method of moments, maximum likelihood, or Bayesian estimators [16, 27, 11, 3]. Several recent methods make no such assumptions and are based on linear programming [39, 37], link Hill numbers to rarefaction/extrapolation sampling curves (richness corresponds to Hill numbers of order 0) [24], perform curve fitting on the rarefaction curve [14, 28] or on ratios of consecutive frequency counts [42]. A recent comparative review by Schmitz and Rahmann [36] shows that despite this long history, there remains an unmet demand for accurate richness estimators when the population is more than ten times larger than the observed sample.

For entropy, classical asymptotic estimators include the Miller-Madow estimator [30] and the coverage-adjusted estimator (CAE) [10], complemented by more recent linear-programming-based estimators [39]. Because Hill numbers of order 1 correspond to the exponential of the Shannon diversity, iNext [24] can also be used to extrapolate entropy. For a broader overview of entropy estimators, we refer the reader to the survey by Paninski [33]. Two practical issues complicate richness (and entropy) estimation in DNA sequencing applications. First, sample diversity (in particular, sample richness) is highly correlated with sequencing depth, which has led to the widespread practice of comparing samples after rarefaction [15, 35]. However, rarefaction discards information, so extrapolating estimators are preferable in many settings [13, 9, 25]. Second, sequencing errors inflate the singleton count, leading to biased estimates. Many standard error correction methods remove all singletons, which violates the assumptions of estimators such as Chao 1 [12, 6]. These two issues increase the demand for estimators that are less dependent on the sequencing depth and the number of singletons.

### The uncertainty quantification gap

A diversity estimate is only as useful as the uncertainty attached to it, since many downstream questions are comparison tasks for which a point estimate alone does not yield valid conclusions. Yet uncertainty quantification has received much less attention than point-estimate accuracy in the above mentioned literature. Several widely used entropy estimators (Miller-Madow, CAE, Unseen) report no confidence intervals at all. Among extrapolating estimators that do, iNext provides analytic intervals, PreSeq provides bootstrap intervals, and RichnEst provides intervals derived from a per-term linear proxy, but to our knowledge none has been systematically shown to achieve nominal coverage in the regime where the population is more than ten times larger than the observed sample. We show in Section 3 that this gap is sharp: across six simulated distribution families and two biological datasets, the nominal 95% intervals of iNext, PreSeq, and RichnEst miss the truth in up to 80% of instances, well above the nominal 5%.

### Contributions

We close this gap with DivQuant, an extrapolating richness and entropy estimator whose 95% confidence intervals empirically cover the truth in essentially all tested regimes. The core change is methodological: we replace the linear objective of DivQuant’s predecessor RichnEst [37], which searches via a linear program for the most plausible population histogram consistent with the observed sample under hypergeometric subsampling, with a quadratic Neyman *χ*^2^ objective [32], turning the search into a convex quadratic program whose statistic is asymptotically *χ*^2^ distributed under certain assumptions. Minimizing and maximizing the population richness over the set of fingerprints whose *χ*^2^ statistic falls below the desired quantile yields confidence intervals. We contribute two supplementary enhancements.

- We replace RichnEst’s fixed-threshold sample truncation with the rare/abundant fingerprint split of Valiant and Valiant [39]. The fixed threshold collapses the degrees of freedom of the *χ*^2^ statistic and frequently renders the confidence-interval program infeasible; the rare/abundant split preserves enough degrees of freedom for the program to remain feasible across a much wider range of inputs.
- We plug the optimal population fingerprint returned by the program into Shannon’s entropy formula to obtain a point estimate, and reuse the same min/max-over-feasible-set construction to obtain entropy confidence intervals. The resulting point estimates outperform classical asymptotic estimators (Miller-Madow, CAE, Unseen) and the extrapolating iNext estimator on both simulated and real data, and the entropy intervals cover the truth in a larger proportion of cases than iNext’s.

In the next section, we describe DivQuant in more detail. In Section 3, we compare DivQuant with state-of-the-art estimators for both richness estimation and entropy estimation on simulated and biological data. We give concluding remarks in Section 4.

## 2 Methods

After introducing the notation, we describe our assumptions and explain how to formulate the estimation problem as a convex optimization problem.

### 2.1 Notation

We are interested in a population with *N* elements (sometimes called individuals) but have only access to a small random sample *x* = (*x*_1_, *x*_2_, …, *x*_*n*_) of size *n < N*. Each element belongs to a single species (sometimes called class or group), and the richness *S* of a population is given by the number of distinct elements (or species).

Assuming that the species have relative frequencies or probabilities *p* = (*p*_1_, *p*_2_, …, *p*_*S*_) in the population, we can compute the (Shannon) entropy by

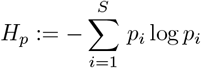

where, depending on the research area, it is common to use log_2_, log_10_ or log_*e*_.

Both species richness and entropy are invariant under relabeling; we will thus in the following consider the *fingerprint* of a sample (sometimes called *frequency* or *occupancy* vector). Given a sample *x* = (*x*_1_, *x*_2_, …, *x*_*n*_), the *k*-th component of the fingerprint *f* = (*f*_1_, *f*_2_, …) counts how many species occurred exactly *k* times in the sample. Note that *f*_0_ corresponds to the unseen species. Thus, the sample size satisfies

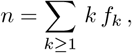

and the observed richness is given by

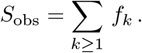

Given the fingerprint, the (empirical) sample entropy can be expressed as

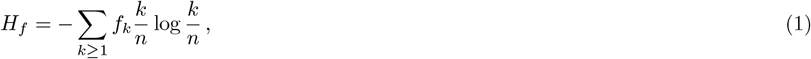

where each species’ probability is approximated by its observed frequency *k/n*. The symbols used throughout the paper are collected in Table 1.

**Table 1.** Notation.

| Symbol | Meaning |
| --- | --- |
| $N, n$ | population size and sample size ($n < N$) |
| $S, S_{\text{obs}}$ | population richness and observed sample richness |
| $H$ | Shannon entropy |
| $x = (x_1, \dots, x_n)$ | sample (one entry per element) |
| $f = (f_0, f_1, \dots, f_{K_{\text{obs}}})$ | sample fingerprint: $f_k$ is the number of species observed exactly $k$ times |
| $K_{\text{obs}}$ | largest count actually observed in $f$ |
| $a = (a_1, \dots, a_C)$ | population fingerprint: $a_c$ is the number of species with abundance $c$ |
| $f' = (f'_0, f'_1, \dots, f'_K)$ | expected sample fingerprint under hypergeometric subsampling from $a$ |
| $C, K$ | truncation parameters for $a$ and $f'$ (Section 2.5) |
| $P_{c,k}^{N,n}$ | hypergeometric probability that a species with abundance $c$ is observed exactly $k$ times when downsampling from $N$ to $n$ elements (Eq. 2) |

### 2.2 Sampling process

We adopt the same probabilistic model as RichnEst [37].

The observed fingerprint *f* is assumed to arise from drawing a random sample of size *n* from the population. Thus, our goal is to find a population of size *N* (or in practice, a larger sample of interest with size *N*), whose *expected* fingerprint *f* ^*′*^ is close to the *observed* fingerprint *f*.

A species that occurs *c* times in the population can be observed 0≤ *k* ≤ *c* times in the sample. The probability that a species with *c* individuals in a population of size *N* is observed exactly *k* times in a sample of size *n* is given by the hypergeometric probability

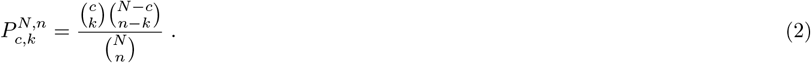

Given a population fingerprint *a* = (*a*_1_, *a*_2_, …, *a*_*C*_), where the most frequent species has *C* individuals and 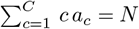, the expected fingerprint 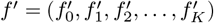 of a sample of size *n* is

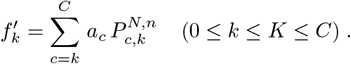

See Figure 1D for an example. We exploit below that the contribution to 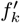 from a species with abundance *c*, namely 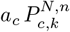, is the mean of a Binomial(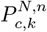) random variable. When the per-trial success probabilities 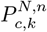 are small, the sum of Binomial random variables is well approximated by a Poisson distribution due to Le Cam’s theorem [29, 23].

### 2.3 Estimation problem

We estimate the species richness by modeling the inverse sampling process to find the most plausible population fingerprint *a* that can explain the observed sample fingerprint *f* = (*f*_1_, *f*_2_, …). The richness estimate is then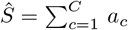.

We formulate the problem as a convex optimization problem and turn our goal that the expected fingerprint *f* ^*′*^ should be close to the observed fingerprint *f* into the objective function. We adopt the minimum *χ*^2^ method [32, 4], which is widely used for parameter estimation in counting experiments whose histogram entries are approximately Poisson distributed [22, 26] and for comparing histograms [2, 17]. Concretely, we use Neyman’s *χ*^2^ statistic [32]

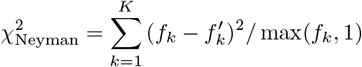

which is asymptotically *χ*^2^ distributed. Following common practice, we use max(*f*_*k*_, 1) in the denominator to handle zero counts [22]. Compared to Pearson’s *χ*^2^ (which uses 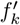 in the denominator), Neyman’s variant has the advantage of yielding a quadratic objective in the variables, hence preserving convexity of the program below. In a good solution, 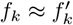, and the difference is marginal; see also Sec. 2.4.2.

Given the observed sample fingerprint *f* = (*f*_1_, *f*_2_, …) of size *n* and the population size *N*, we solve the following convex optimization problem to search for the population fingerprint *a* = (*a*_1_, *a*_2_, …, *a*_*C*_), where 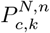 are the hypergeometric probabilities of Equation (2) and the non-negative vector 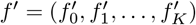 are auxiliary variables that store the expected fingerprint.
minimize 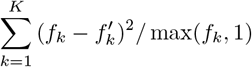

subject to

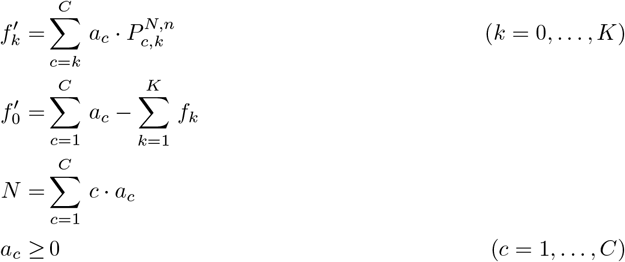

The important variables are the components of the population fingerprint *a*, which determine *f* ^*′*^, which has to be close to the observed *f*. The optimization problem formulation depends on the truncation parameters *C* and *K*, which we do not know initially. We describe their choice in Section 2.5. The constraint 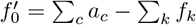 implements 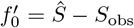 that is, 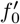 counts the species present in the population but unseen in the sample. We relax both *a* and the auxiliary variables *f* ^*′*^ to be non-negative real numbers rather than integers. While this does not yield strictly valid fingerprints, it makes the program more efficient while empirically leaving the richness estimate essentially unchanged. There may exist several solutions for *a* with (almost) equivalent objective values; we come back to this below. The original formulation of RichnEst uses the linear proxy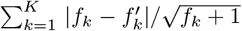, which in practice gives similar point estimates but makes it hard to derive confidence intervals [37].

### 2.4 Confidence intervals

Since many population fingerprints *a* can yield comparable objective values, a confidence interval for the richness estimate is more informative than a single point estimate. We construct the confidence interval by inverting the *χ*^2^ test.

#### 2.4.1 Inverted *χ*^2^ test

The general idea of inverting a *χ*^2^ test is to collect all parameter values under which the hypothesis would not be rejected [1].

For each candidate *a*, the null hypothesis is that *a* is the true population fingerprint. Thus, inverting the tests gives a confidence region for the true *a*. Let 𝒜 be the set of valid population fingerprints whose Neyman *χ*^2^ statistic, computed between their expected fingerprint *f* ^*′*^ and the observed fingerprint *f*, is below the critical value *ε* at the desired confidence level. The minimum and maximum of 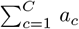 over *a ∈ 𝒜* then form a confidence interval for the true richness *S*.

We therefore minimize or maximize the richness under the constraint that the deviation between the observed and expected fingerprint is bounded by *ε*, whose choice is discussed below. For the lower bound, we thus solve

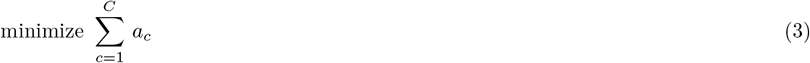

subject to

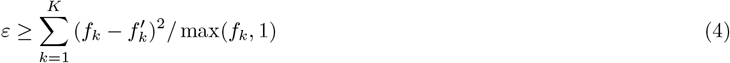

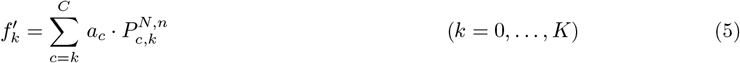

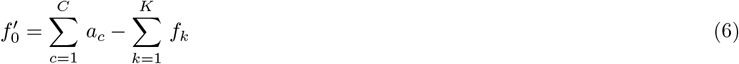

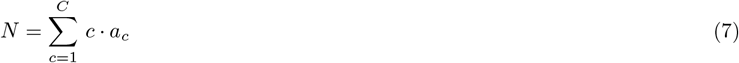

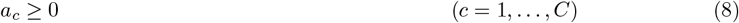

and, respectively, maximize objective (3) to get the upper bound.

We set *ε* to the value of the quantile function of a *χ*^2^ distribution with *K −* 1 degrees of freedom for the desired confidence level (default 0.95). This is a heuristic choice, but one for which there are good reasons. Under the null hypothesis, by Le Cam’s theorem, the (*f*_*k*_)_1*≤k≤K*_ are approximately independent Poisson variables [29] linked by the single constraint 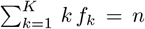 Thus, Neyman’s *χ*^2^ statistic asymptotically follows a 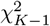 distribution. The unseen species 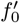 do not enter the *χ*^2^ sum and hence do not affect the degrees of freedom. Although the optimization later searches over *a*, the test inversion fixes *a* when evaluating the statistic: for each candidate *a* we ask whether the observed *f* is compatible with *a* as a fixed null. Because *a* is not estimated from *f*, the usual degrees-of-freedom penalty for fitted parameters does not apply.

#### 2.4.2 Limitations of the *χ*^2^ approximation

The *χ*^2^ approximation only holds under several assumptions, which are partially violated or hold only asymptotically in our setting. The main limitations are the following:

▃ The standard rule is that the *expected* counts 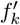 should be at least 5. This assumption is not always true in our setting, particularly in the tail, where 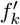 may be small.
▃ Agresti and Ryu [1] described the test inversion for Pearson’s *χ*^2^ statistic rather than Neyman’s. We use Neyman’s *χ*^2^ statistic for the convexity reason described in Section 2.3.
▃ The program is feasible if and only if the minimum of Neyman’s *χ*^2^ statistic (obtained by the optimization problem in Section 2.3) is at most *ε*. Since *ε* grows with the degrees of freedom, feasibility is sensitive to the choice of *K*. However, the value of *K* is not uniquely determined and can be increased by appending up to *C − K* zero entries to the observed fingerprint. By using the ideas of Section 2.5 and Section 2.6, we obtain feasible optimization problems in practice.

Due to these limitations, the confidence interval construction is best understood as a heuristic rather than a fully derived asymptotic procedure. We show in Section 3 that the resulting confidence intervals empirically achieve near-nominal coverage. The empirical tendency toward over-coverage rather than under-coverage may be explained by two conservative effects: When the expected counts 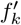 are small, the distribution of 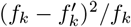 is right-skewed compared to its asymptotic Gaussian or *χ*^2^ approximation, so a symmetric *χ*^2^ critical value inflates *ε*; and using max(*f*_*k*_, 1) in the denominator further tames the contribution of small counts.

### 2.5 Estimating truncation parameters *C* and *K*

The explicit modeling of the sampling process depends on the parameters *C ≤ N* and *K ≤ n*, which determine the length of the variable vectors *a* and *f* ^*′*^. We follow the strategy of Schröder and Rahmann [37] to determine *C* and *K*. We determine them in sequence: *K*_obs_ is read from *f*, then *C* is chosen from *K*_obs_ via a threshold *t*, then *K* is chosen from *C* via a threshold *τ*.

Let *K*_obs_ denote the largest count observed in the sample fingerprint *f*. Assuming that we have a species with abundance *c* in the population, the probability to observe a count larger than *K*_obs_ in our sample is given by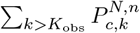. Since no such count was observed, we require that the tail probability 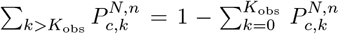 is small. Given a threshold *t ∈* (0, 1), we therefore set

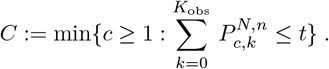

Lower values of *t* are more conservative and lead to a larger *C*. This creates a trade-off between being conservative and the size of the program.

Since the optimization program only measures the deviation between the observed and expected fingerprint up to *K*, we must ensure that essentially all the probability mass lies in the considered range. For example, if we observe no species with more than 10 individuals, the entries in 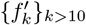 should also be near zero. For a given *C* and a threshold *τ ∈* (0, 1), we set

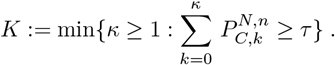

The observed fingerprint is padded with *K− K*_obs_ trailing zeros.

The two thresholds play different roles: *t* caps *C* from above, so the more conservative (smaller) *t*, the larger *C* and the more expensive the program; *τ* caps how much tail mass of the per-*c* subsampling distribution we allow to be ignored, with little additional cost when set close to 1. We therefore use *t* = 0.8 and *τ* = 0.99 as default parameters. The resulting richness estimates are insensitive to the choice of *t* and *τ* as long as both parameters are within a reasonable range (see Appendix B).

### 2.6 Reducing the problem size

A challenge arises for samples where both *K* and *C* are large, resulting in impractical solving times that exceed the typical few seconds, see Appendix A. RichnEst reduces the program size by truncating the observed fingerprint at a fixed threshold *K*_max_ (default 25), only retaining 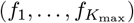 and rescaling *n* and *N* accordingly [37]. However, this threshold is not very robust to varying input properties. A small threshold reduces the degrees of freedom of the *χ*^2^ distribution, which can render the confidence-interval program infeasible. The extent to which this affects the solvability of the confidence interval is shown in Appendix A. We instead use the rare/abundant fingerprint split proposed by Valiant and Valiant [39], which avoids the choice of *K*_max_. If a species is very abundant in the population — say, it makes up half — then its sample frequency will be close to its expected value *n/*2. Thus, we can use the trivial empirical estimate of *N/*2 for this species.

Concretely, an abundant species’ sample frequency is tightly concentrated around its expectation, so in the fingerprint *f* the values *f*_*i*_ for *i* near that frequency are small; only a few species contribute counts in that neighbourhood. We therefore say that a frequency *k* is *abundant* if its neighbourhood in *f* is sparse:

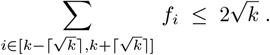

We remove each *abundant f*_*k*_ from the fingerprint, give it an estimated population abundance of *k ·N/n*, and reduce *n* and *N* by *k·f*_*k*_ and *k ·f*_*k*_*·N/n*, respectively. After solving the reduced program, the empirical entries are added back to recover the full population fingerprint *a*.

This split has been shown to work well in practice [39]. The plug-in is justified because the sparse-neighbourhood criterion identifies precisely those frequencies whose corresponding species are tightly concentrated around their expectation by Le Cam’s theorem, so *k ·N/n* is itself a low-variance estimator and the corresponding entropy contribution *−*(*k/n*) log(*k/n*) closely tracks the population value *−*(*c/N*) log(*c/N*).

### 2.7 Entropy estimation

The optimization problem directly yields an estimate of the population fingerprint *a*. We thus estimate the entropy of a population with upsampled fingerprint *a* = (*a*_1_, *a*_2_, …, *a*_*C*_) and population size *N* as

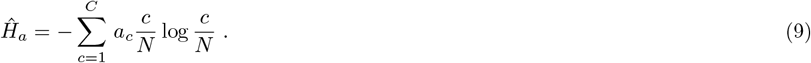

That is, we plug the estimated population fingerprint instead of the observed sample fingerprint into Equation (1).

To obtain entropy confidence intervals, we replace Equation (3) by Equation (9) and minimize or maximize the optimization program from Section 2.4.1 to obtain the lower and upper bound of the confidence interval, respectively.

## 3 Results

We compare DivQuant against the observed richness *S*_obs_ and the best-performing extrapolating estimators from the review by Schmitz and Rahmann [36] on simulated and biological data, namely, its predecessor, RichnEst [37] (https://gitlab.com/rahmannlab/richnest, commit de571d68), iNext [24] (R package, version 3.0.2), and PreSeq [14] (R package, version 4.0.0).

For entropy estimation, we compare DivQuant against the observed sample entropy, given by Equation (1), the asymptotic Miller-Madow bias-corrected estimator [30], the coverage adjusted estimator (CAE) [10], Unseen [39] and the extrapolating entropy estimator available in iNext [24]. For Unseen, we use our Python re-implementation of the original MATLAB code. DivQuant, RichnEst and our re-implementation of Unseen use the Gurobi Python API to solve the (linear or quadratic) convex optimization problems [21]. All experiments were run on an Ubuntu (24.04 LTS) server with two AMD EPYC 9534 64-Core Processors, 1.5 TB of 4800-MHz DDR5 memory, and a KIOXIA CD8P SSD. We measured running times and maximum memory usage with /usr/bin/time -v.

### 3.1 Simulated Data

We simulate data with different species compositions using the statistical population models described in the review by Schmitz and Rahmann [36]. We set the population size to *N* = 10^5^ and the richness to *S* = 10^4^. Given a simulated population, we create samples that contain 1% to 90% of the population. We denote the absolute abundance vector of a population by *X* = (*X*_1_, …, *X*_*S*_), and use *α* to denote the appropriate scaling factor such

That 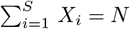. The simulated species composition models are as follows.

In the *Random* model, we draw *p*_*i*_ for *i* = 1, …, *S* from the uniform distribution on [0.02, 0.98] and set *X*_*i*_ = *α ·p*_*i*_.

In the *Homogeneous* model, every species *i* = 1, …, *S* has the same abundance of *N/S* (here 10^5^*/*10^4^ = 10).

In the *Negative Binomial* model, we draw a random sample *X*^*′*^ of size *S* from a Negative Binomial distribution either with parameters *r* = 2 and *p* = 0.02 (model 1), or with *r* = 20 and *p* = 0.2 (model 2). For *i* = 1, …, *S*, we set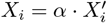.

To create a bimodal composition (*Two Mixture* model), we draw a random sample *X*^*′*^ from a Negative Binomial distribution with parameters *r* = 2 and *p* = 0.1 for one half of the species and with parameters *r* = 50 and *p* = 0.2 for the other half of the species. The absolute abundances are then given according to the description above.

In the *Geometric* model, we draw a sample *X*^*′*^ from a geometric distribution with parameter *p* = 1*/S* and set 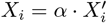 such that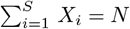.

In the *Zipf-Mandelbrot* model, *X*_*i*_ = *α/*(*i* + 10) for all *i* = 1, …, *S*.

#### 3.1.1 Richness estimation

We first compare the richness estimates of DivQuant, RichnEst, iNext, and PreSeq on simulated data. Figure 2 shows the log_2_ ratio between the estimated and true richness, aggregated across all distribution types.

**Figure 2.**
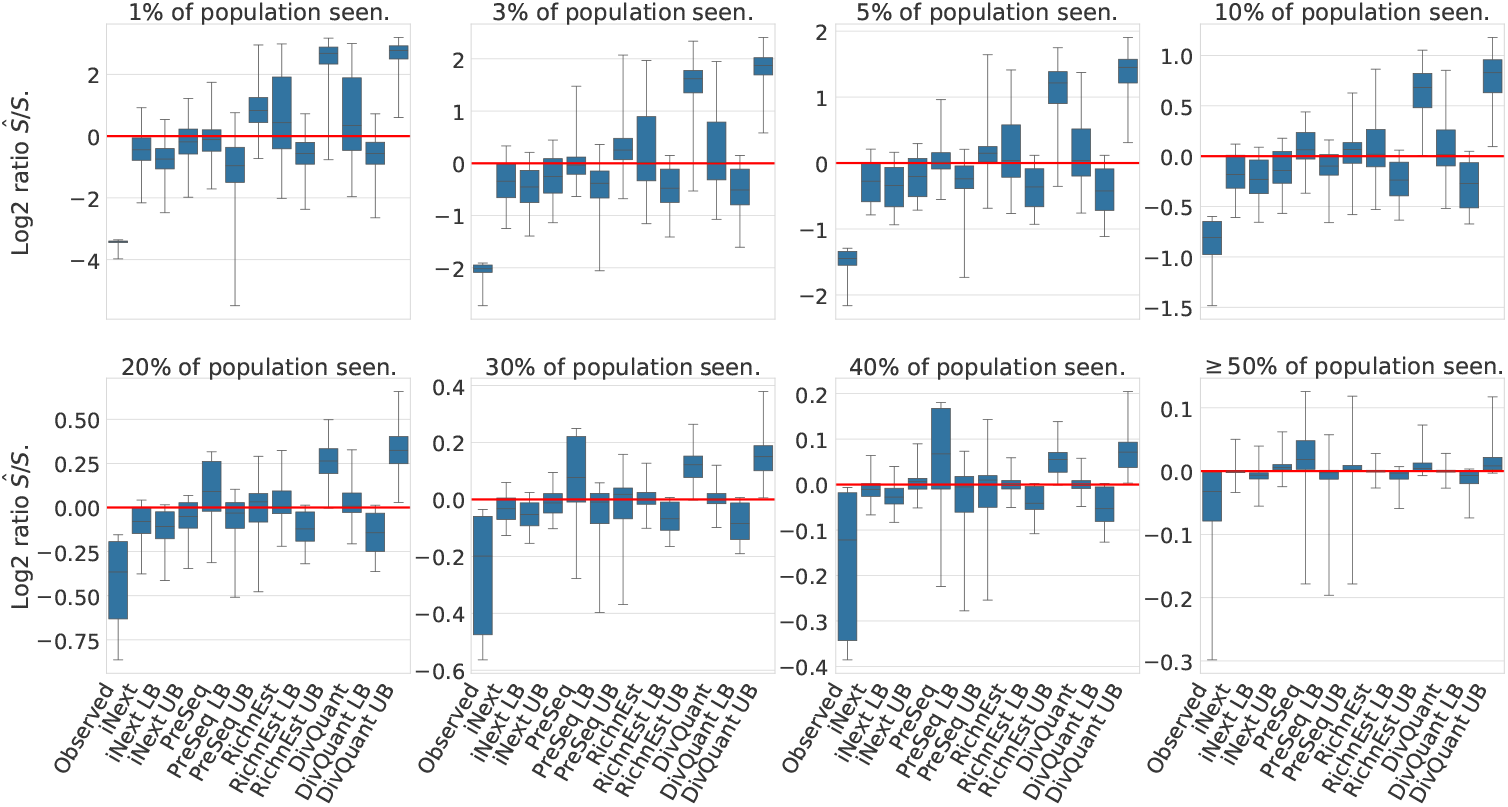
Log_2_ ratios between estimated richness and population richness across different samples from the simulated data. A log_2_ ratio of zero corresponds to a perfect correspondence, while log_2_ ratios *<* 0 underestimate the true richness and log_2_ ratios *>* 0 overestimate the true richness. Each method’s point estimate, together with the lower and upper bounds of the computed 95% confidence interval, is shown. DivQuant’s intervals are wider than RichnEst’s by design; Figure 3 shows that this width translates into well-calibrated coverage, whereas the narrower intervals of competing methods systematically miss the truth.

As expected, the observed sample richness underestimates the true population richness, and all extrapolating estimators improve as the sampling fraction grows. For small samples (*<* 10%), PreSeq yields more accurate point estimates compared to iNext, RichnEst, and

DivQuant. For larger samples (*>* 10%), both RichnEst and DivQuant are more accurate than PreSeq and iNext. The point estimates of RichnEst and DivQuant are similar throughout, but DivQuant’s confidence intervals are wider.

While narrower 95% confidence intervals are generally preferred, an interval is only meaningful if it covers the true value at the nominal rate. Neither iNext nor PreSeq achieves this. The upper bound of iNext is often below the true richness, and PreSeq’s bootstrap intervals frequently fail to cover the point estimate.

This is quantified in Figure 3. DivQuant’s intervals contain the true richness in essentially all instances, except for the Homogeneous and Negative Binomial models at small sampling fractions (*≤* 3%). For sampling fractions above 50%, DivQuant’s intervals are well-calibrated, whereas more than 5% of RichnEst’s intervals fail to cover the true richness. For small sampling fractions, fewer than 20% of iNext intervals and at most 80% of PreSeq intervals cover the true richness. Even at a sampling fraction of 40%, only about half of iNext’s intervals and roughly 20% of PreSeq’s intervals are correct.

**Figure 3.**
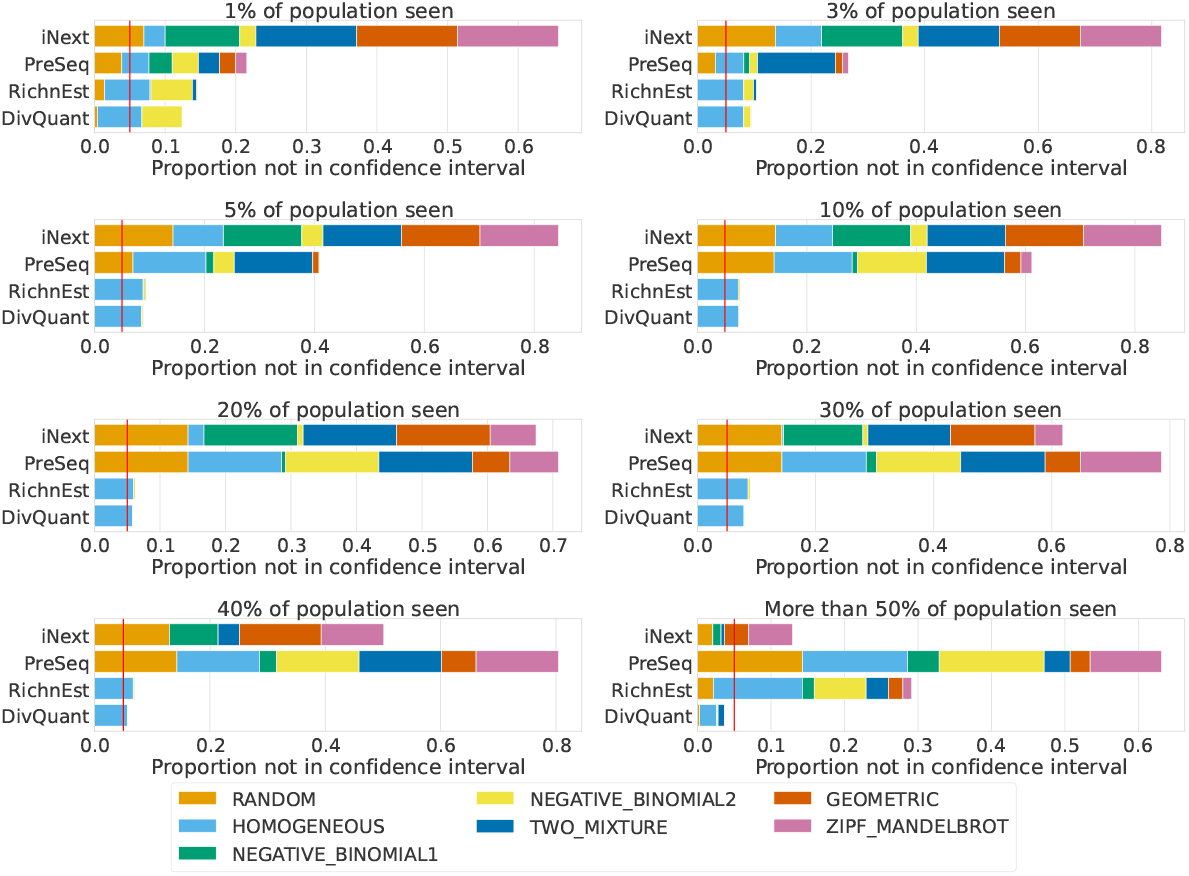
Fraction of simulated instances where the true richness is not in the 95% confidence interval; split for different sampling fractions. The vertical red line marks the theoretically allowed portion of instances (5%) that are not within the confidence interval. Colors indicate the distribution model from which the failed instances originate.

In summary, of the four methods compared, DivQuant is the only one whose confidence intervals closely approach the nominal 95% level across the simulated samples. By contrast, iNext’s and PreSeq’s confidence intervals seldom contain the true richness.

#### 3.1.2 Entropy estimation

We next compare the estimated entropy on simulated data. Since iNext computes entropy in base 10, we report all entropies using log_10_. DivQuant supports log_2_, log_10_ and log_*e*_.

Figure 4 shows that the observed sample entropy generally underestimates the true entropy. For larger sampling fractions, the asymptotic estimators (Miller-Madow, CAE, Unseen), and the extrapolating estimator, iNext, begin to overestimate the true entropy. DivQuant yields the most accurate entropy estimates for sampling fractions *>* 3%.

**Figure 4.**
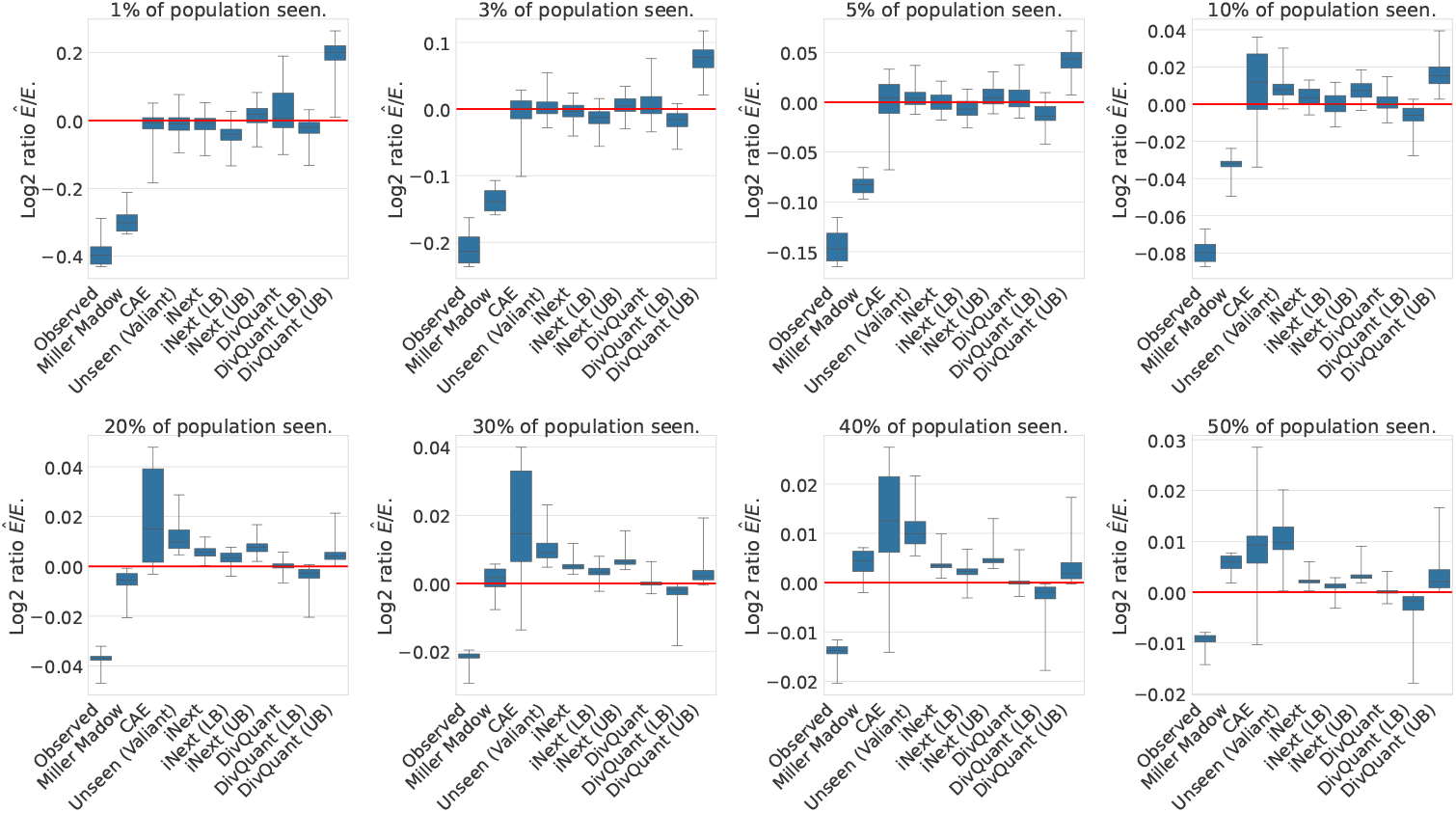
Log_2_ ratio between estimated entropy and population entropy across different samples of the simulated data. For iNext and DivQuant, LB and UB denote the confidence intervals. A log_2_ ratio of zero corresponds to a perfect correspondence, while log_2_ ratios *<* 0 underestimate the true entropy and log_2_ ratios *>* 0 overestimate the true entropy.

Miller-Madow, CAE and Unseen do not provide confidence intervals. The confidence intervals of iNext do not contain the true entropy for a large portion of tested instances, see Figure 4 and Figure 5. By contrast, DivQuant’s confidence intervals contain the true entropy in a larger proportion of cases. For a small sampling fraction, more than 5% of instances are not within the confidence interval; for a larger sampling fraction, only estimates for the homogeneous population are not within the confidence interval.

**Figure 5.**
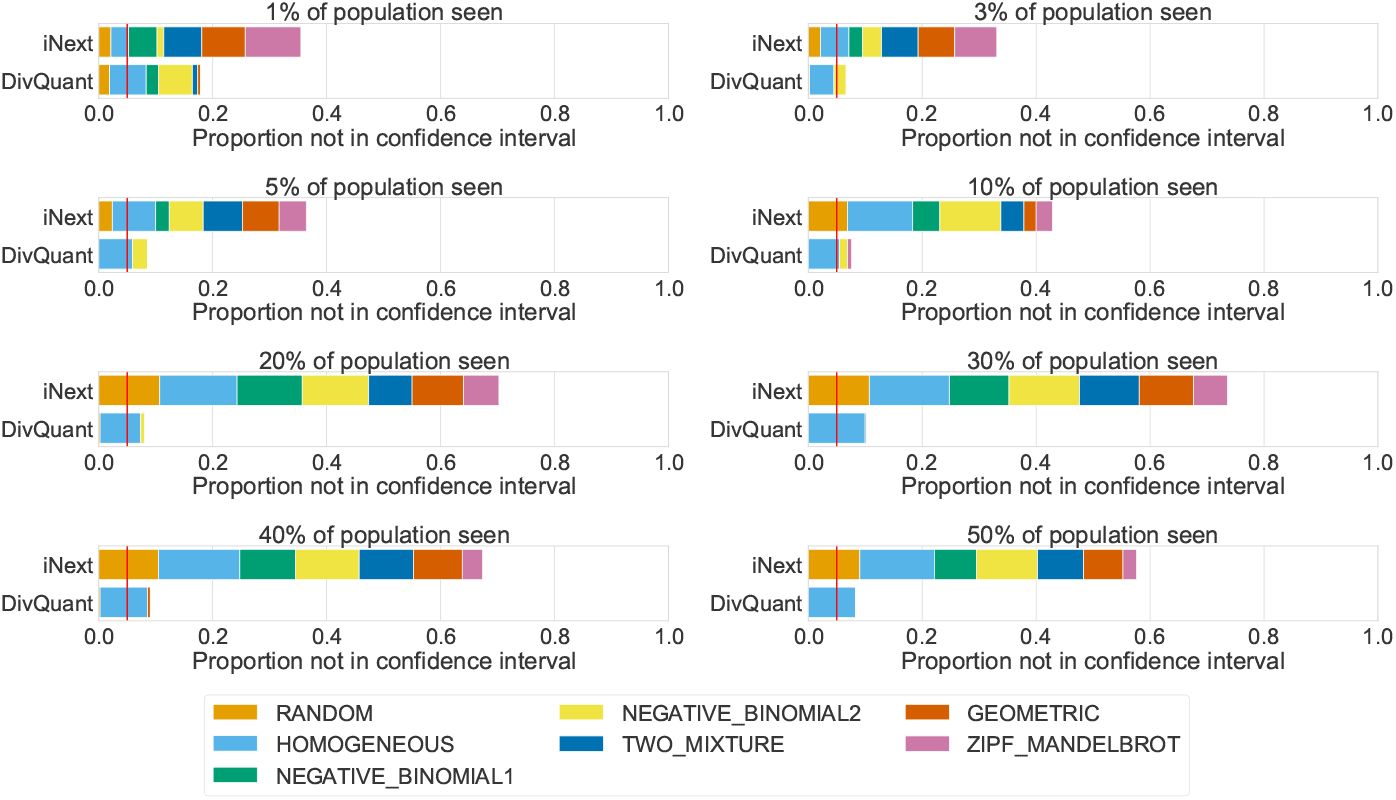
Fraction of simulated instances where the true entropy is not in the 95% confidence interval; split for different sampling fractions. The vertical red line marks the theoretically allowed portion of instances (5%) that are not within the confidence interval. Colors indicate the distribution model from which the failed instances originate.

### 3.2 Microbiome richness

We obtained the taxonomic profiles from the *Tara Oceans* consortium [5], which sequenced plankton samples across all major oceanic regions. From the operational taxonomic unit (OTU) count tables of the 139 different samples (here treated as the “population”, available at ocean-microbiome.embl.de/companion.html), we drew random subsamples that contain *ρ* = 0.01, 0.03, 0.05, 0.1, 0.2, 0.3, 0.4, 0.5 of the complete sample. For each sample (population), we compared the estimated richness on the subsample with the richness of the complete sample. A fair comparison with asymptotic estimators is not possible because the full underlying population is unknown. We nevertheless report entropy results for completeness in Appendix C.

Figure 6 shows that, similarly to the results on simulated data, all methods yield more accurate richness estimates and smaller confidence intervals as the sampling fraction increases. DivQuant has larger confidence intervals compared to PreSeq and RichnEst. However, both PreSeq and RichnEst have a larger portion of instances (6% and 10%) where the correct richness is not contained within the confidence interval (see Figure 7A). The upper bound of iNext underestimates the true richness in more than half of the instances.

**Figure 6.**
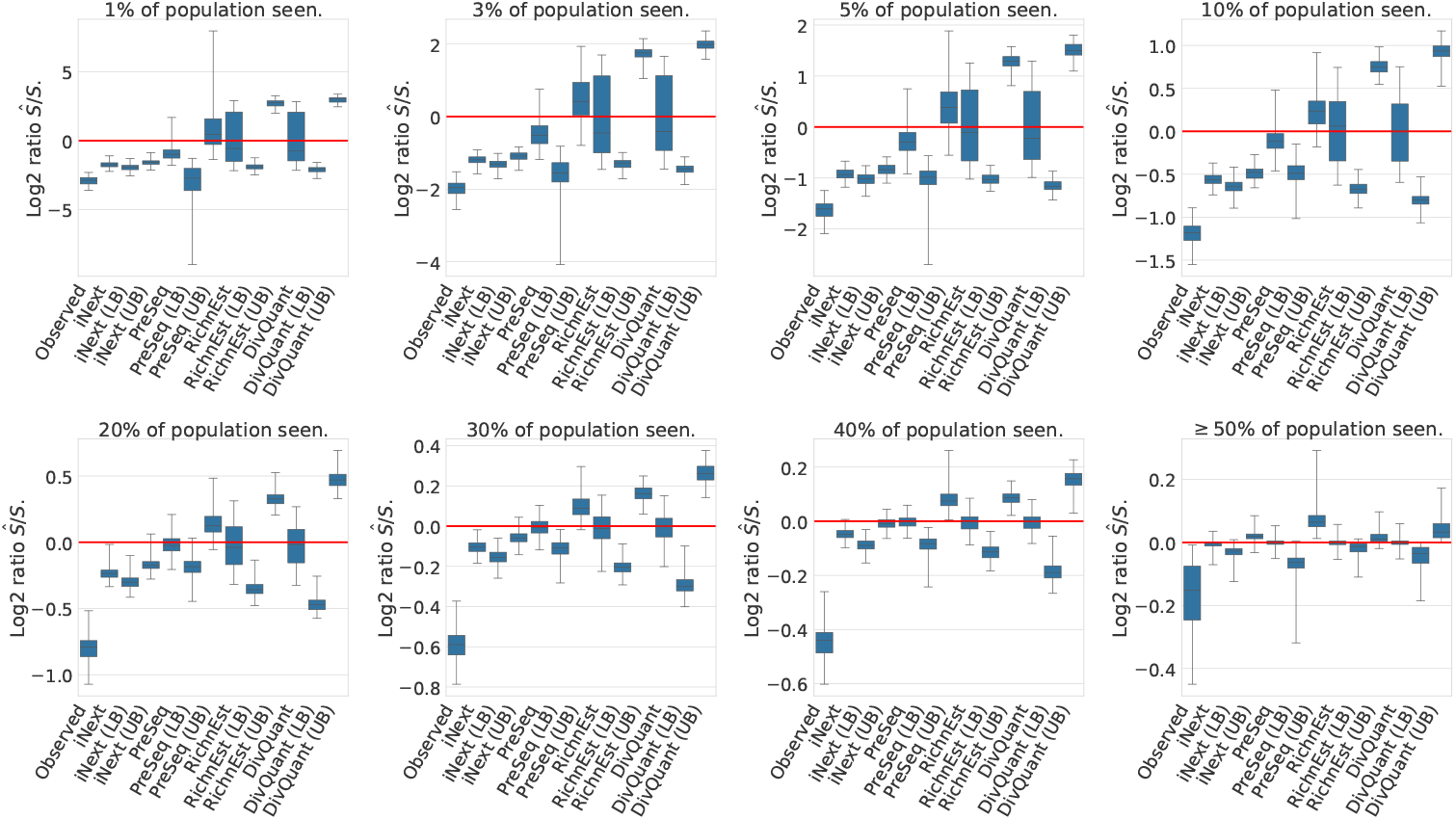
Log_2_ ratio between estimated richness and population richness for the *Tara Oceans* microbial taxonomic profiles across different sampling fractions. A log_2_ ratio of zero corresponds to a perfect correspondence, while log_2_ ratios *<* 0 underestimate the true richness and log_2_ ratios *>* 0 overestimate the true richness. For all methods, the point estimate and the lower and upper bounds of the computed 95% confidence interval are shown.

**Figure 7.**
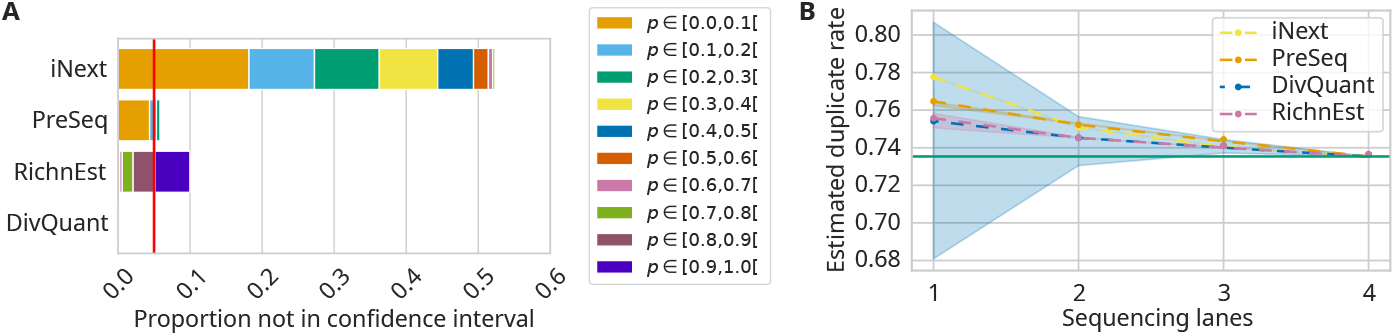
**A** Fraction of *Tara Oceans* instances where the true richness is not in the 95% confidence interval. Colors indicate the sampling fraction. **B** Confidence interval for estimated duplicate rates for the *10X Genomics* scRNA-seq data. Lanes 1 to 4 cumulatively contain *n* = 96 039 314, 192 665 228, 288 186 226, and 382 950 653 sequences, with observed richnesses *S*_obs_ = 54 405 046, 77 172 440, 90 772 183, 101 353 538, respectively. The horizontal green line corresponds to the duplicate rate in the union of all 4 sequencing lanes.

### 3.3 Duplicate rate estimation

We downloaded the scRNA-seq data generated by *10X Genomics* from https://cg.10xgenomics.com/samples/cell-exp/3.0.2/5k_pbmc_v3/5k_pbmc_v3_fastqs.tar. Each read in this data set contains a sequence tag of length 28 that should be unique for each distinct molecule. Given the number *S* of distinct sequence tags in an experiment with *N* reads, the duplicate rate is (*N− S*)*/N*. In practice, one is often interested in the duplicate rate at an increased sequencing depth (for example, sequencing 2*×* to 10*×* as many reads) in order to determine whether further sequencing would result in new molecules being sequenced. We counted the sequence tags using the *k*-mer counter Hackgap [44].

To obtain a ground-truth duplicate rate at increased depth, we pool the reads from all four lanes and use the duplicate rate of the pool (horizontal green line in Figure 7B) as the target. By adding more and more individual lanes to a sample, we obtain samples of increasing size and smaller confidence intervals. The true target value should be inside the confidence interval, and the point estimate should be close to the target.

Figure 7B shows the estimated duplicate rate. Consistent with previous results, DivQuant has the largest confidence intervals but is the only method where the true duplicate rate lies within the confidence interval. All other methods overestimate the duplicate rate, i.e., underestimate the true richness (not shown).

### 3.4 Running time and memory requirements

DivQuant estimates both richness and entropy within a few seconds on both simulated and real data. On all *Tara Oceans* samples, the maximum CPU time is 17 s, and the maximum memory usage is 206 MB across all samples and sampling fractions.

Figure 8 compares the running times of the methods (both richness and entropy estimators) for different subsampling fractions for the `TARA_004_DCM_0.22-1.6` sample. All reported times are averaged over three runs.

**Figure 8.**
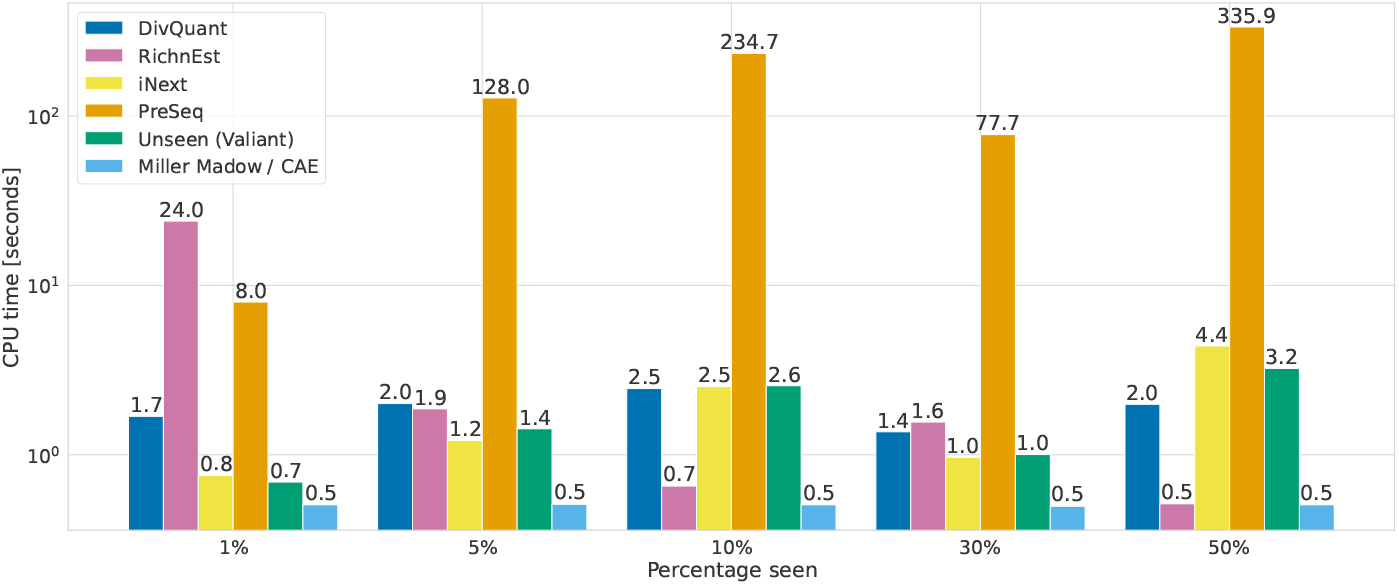
Running time in CPU seconds for the *Tara Oceans* sample TARA_004_DCM_0.22-1.6 (*N* = 143 169). Running times are shown for richness estimators (RichnEst, PreSeq), entropy estimators (Unseen (Valiant), Miller-Madow, CAE), and estimators that estimate both richness and entropy (DivQuant, iNext). The running times (y-axis, log scale) are averaged over three runs for five different sampling fractions (x-axis).

As expected, the Miller-Madow and CAE estimators are fastest, since they have closed-form expressions. On this dataset, DivQuant, iNext, and Unseen need only a few seconds at all subsampling fractions. RichnEst takes only a few seconds for all subsampling fractions except 1%, where it takes 24 s. PreSeq is the slowest method and scales poorly, from about 8 s at 1% to roughly 5 min 36 s at 50%.

## 4 Discussion and conclusion

We have presented an optimization approach to estimate richness and entropy of a population when given access to only a small sample at a known sampling fraction. For richness, DivQuant provides both a point estimate and a confidence interval. For entropy, a point estimate is currently provided.

Our experiments indicate that our new approach computes more accurate richness and entropy confidence intervals compared to other state-of-the-art methods. For sampling fractions *<* 10%, the richness confidence intervals are relatively broad, but the true solution falls within the theoretical 95% confidence interval for nearly all the tested instances. The main exception is the homogeneous distribution: when all species share the same abundance, the sample fingerprint concentrates near a single value *k ≈ n/S*, leaving the *χ*^2^ statistic with essentially one informative bin and the test inversion with little leverage. This affects richness coverage at small sampling fractions and entropy coverage at larger sampling fractions; a tailored treatment of this corner case is left for future work. In contrast, more than 5% of the confidence intervals computed by RichnEst, iNext, and PreSeq fail to cover the true richness. For entropy, plugging the optimal population fingerprint into Shannon’s entropy formula produces more accurate point estimates than the asymptotic estimators (Miller-Madow, CAE, Unseen) and the extrapolating estimator iNext. In addition, the entropy confidence intervals of DivQuant contain the true solution in more instances compared to iNext.

In future work, we aim to further tighten the statistical analysis for richness estimation and to increase the accuracy of entropy confidence intervals.

DivQuant requires the population size *N* to be known. When *N* is only known approximately, DivQuant can be run over a range of plausible *N* values and the minimum lower-bound richness across that range reported as a conservative estimate; because larger *N* adds unseen species rather than restructuring the observed fingerprint, this lower bound tends to saturate. A systematic study of robustness under misspecified *N* is left for future work.

DivQuant currently relies on Gurobi (and its Python API), for which a free academic license is available. Evaluating open-source QP backends (e.g., OSQP, Ipopt, or CVXPY’s SCS/clarabel) is planned as future work.

In summary, DivQuant is well-suited to estimate richness and entropy in the extrapolating setting, where the population size (or a reliable estimate of it) is known.

## 5 Data and code availability

DivQuant and all Snakemake [31] workflows to reproduce the experiments are available at https://gitlab.com/rahmannlab/divquant (commit c005cc33). The optimization problems are solved using a free academic Gurobi license [21].

## APPENDIX

### A Effect of reducing the problem size

We reduce the optimization problem size by splitting the observed fingerprint into two parts. The first part (low abundances) is used for the upsampling process, and the second part (high abundances) is upsampled by multiplying the frequency count by *N/n*.

#### Effect on running time

Reducing the problem size decreases the running time for large samples (Figure 9) for to two reasons. First, we have to compute *C× K* hypergeometric probabilities to compute the expected fingerprints. Second, the optimization problem has more variables and can thus be harder to solve.

**Figure 9.**
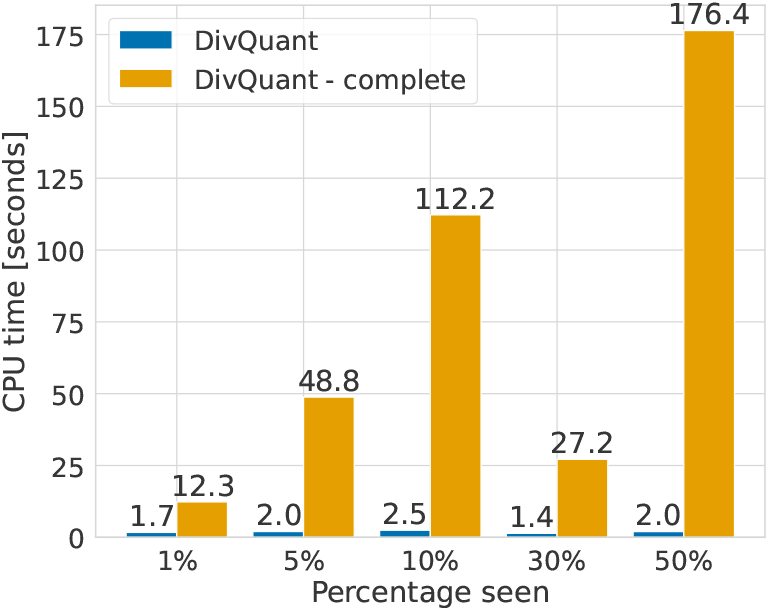
Running time in CPU seconds (y-axis) for different sampling fractions (x-axis) for the *Tara Oceans* sample `TARA_004_DCM_0.22-1.6` (*N* = 143 169). Running times are shown for DivQuant with (standard) and without (complete) reducing the problem size. The running times are averaged over three runs.

#### Effect on richness estimates

Since we only remove abundant species, whose population frequencies are concentrated around their up-scaled sample frequencies, both versions result in similar point estimates, as demonstrated for one microbial sample in Table 2. Since we have a larger *χ*^2^ error bound for larger *K*, the confidence interval is larger using the complete fingerprint compared to using only the rare part of the fingerprint to upsample the population. Hence the wider intervals at small sampling fractions in Figure 2 are intrinsic to the *χ*^2^ inversion when the sample carries little information about the unseen mass, rather than an artefact of the rare/abundant split.

**Table 2.** Richness estimates for DivQuant with (standard) and without (complete) reducing the problem size on the *Tara Oceans* sample TARA_004_DCM_0.22-1.6 (*S* = 4398, *N* = 143 169). LB and UB denote the lower and upper bounds of the 95% confidence interval.

| Estimator | $n/N$ | 1% | 5% | 10% | 30% | 50% |
| --- | --- | --- | --- | --- | --- | --- |
| point |  | 1810 | 7322 | 4476 | 4417 | 4408 |
| point (complete) |  | 1837 | 7388 | 4527 | 4458 | 4438 |
| LB |  | 1189 | 2084 | 2771 | 3751 | 4155 |
| LB (complete) |  | 994 | 1816 | 2312 | 3178 | 3622 |
| UB |  | 41609 | 12664 | 8256 | 5440 | 4634 |
| UB (complete) |  | 53818 | 19706 | 13690 | 7973 | 6126 |

#### Effect on the fraction of solved instances

Figure 10 shows how problem reduction affects the feasibility of the confidence-interval program: Using a fixed threshold (truncation at *K*_max_ = 25, as in RichnEst [37]) leaves substantially more instances without a feasible solution than the rare/abundant split used in DivQuant.

**Figure 10.**
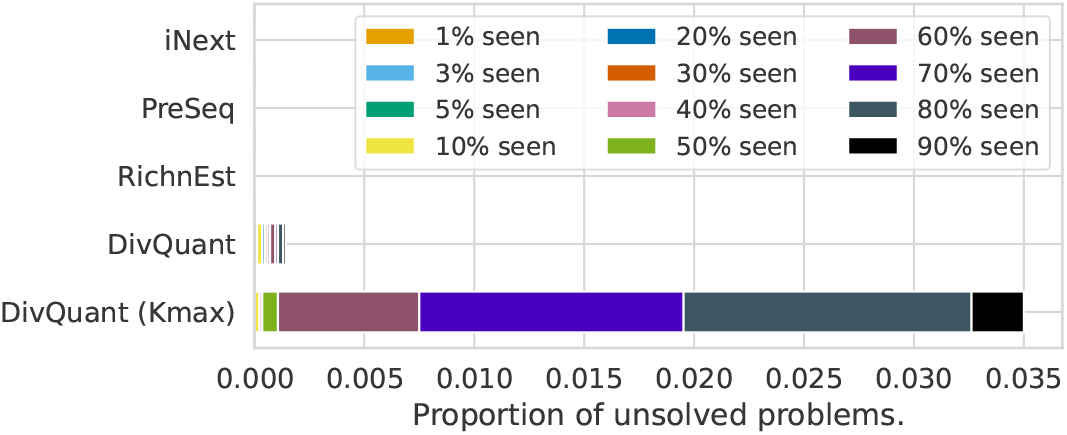
Fraction of simulated instances for which no confidence interval could be computed. DivQuant uses the rare/abundant fingerprint split, while DivQuant (*K*_max_) truncates at the same fixed threshold of 25 as RichnEst. The splitting scheme is more robust and leaves fewer instances without a feasible confidence-interval program.

### B Choosing thresholds *t* and *τ* to estimate the problem size

Figure 11 shows the mean log_2_ ratio between the estimated and true richness for different parameter combinations of *t* and *τ*, which determine *C* and *K*. Figure 12 shows the corresponding standard deviation for the point estimates.

**Figure 11.**
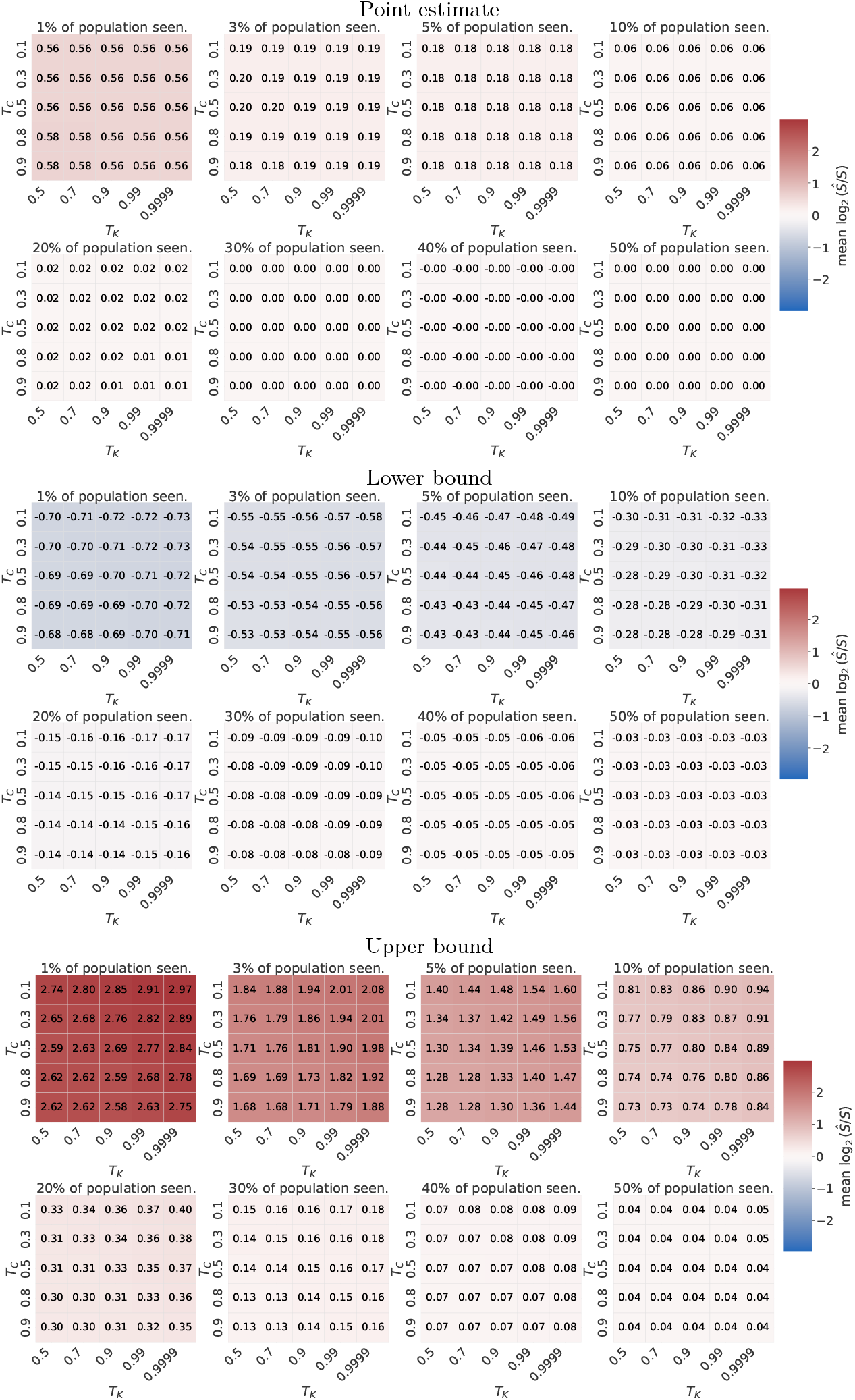
Mean log_2_ ratios of the estimated (point estimate, lower, and upper bounds) and true richness for different parameter combinations of *t* and *τ*. A log_2_ ratio of zero is best.

**Figure 12.**
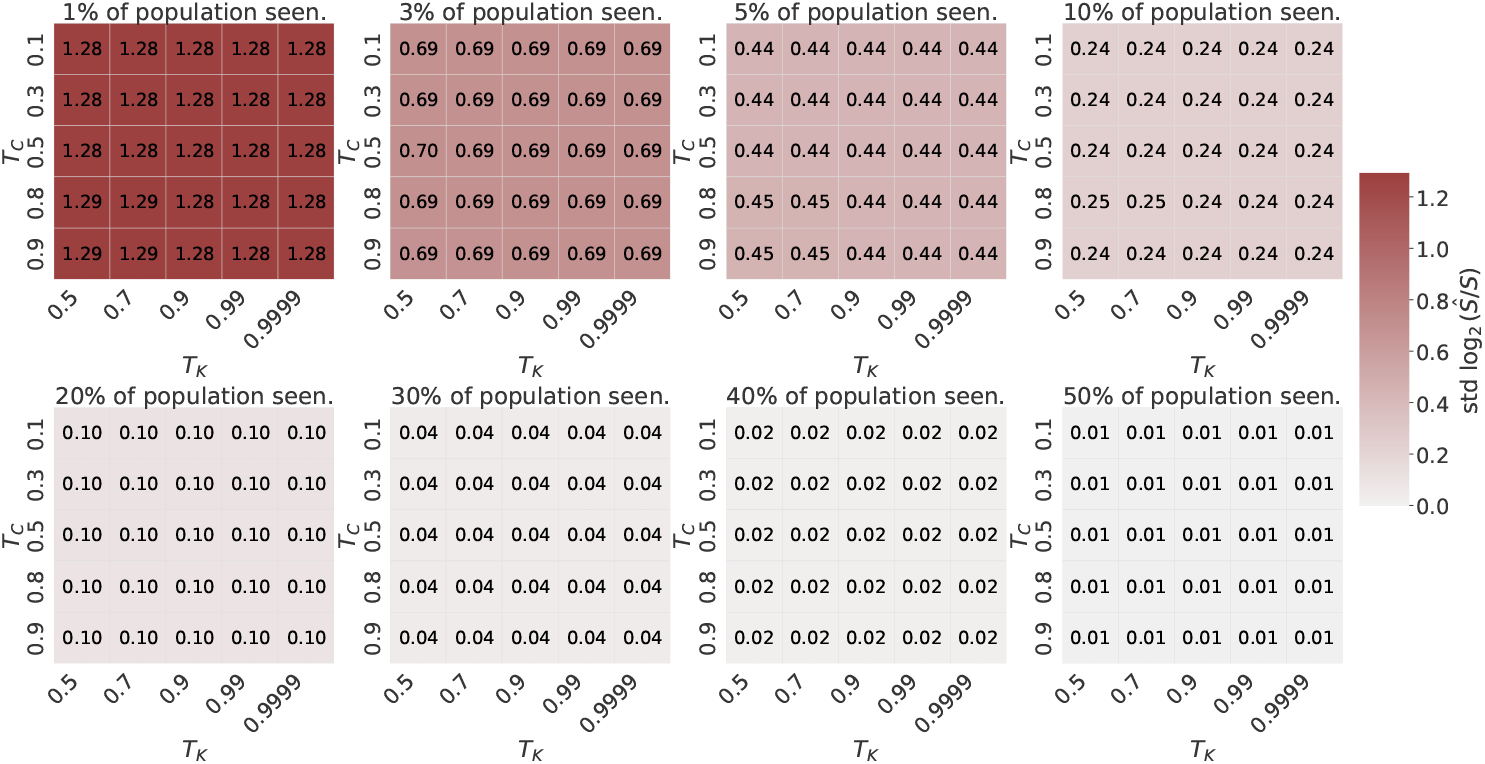
Standard deviation of the log_2_ ratios between the estimated (point estimate, lower and upper bound) and true richness for different parameter combinations of *t* and *τ*. Smaller values (close to zero) are better.

DivQuant’s point estimates have a small positive bias, in particular for small sampling fractions. The variation of the point estimate decreases for increasing sample sizes. As expected, the lower and upper bounds yield negative and positive log_2_ ratios, respectively.

The dependence on *t* and *τ* is negligible within a reasonable parameter range. For example, the point estimates are slightly worse if *t* ≥ 0.8 and simultaneously *τ ≤* 0.7 at the smallest sampling fraction, but for all other combinations, the mean and standard deviation are almost identical.

### C Estimated entropy on the microbiome data

We estimated the entropy for the data discussed in Section 3.2. Since we do not have access to the whole population, we consider the complete sample as the population. Thus, the estimates of the asymptotic estimators (Miller-Madow, CAE, and Unseen) may deviate from the *true* entropy. Since iNext and DivQuant are extrapolating estimators, their entropy estimates should be close to the *true* entropy.

The entropy estimates are shown in Figure 13. For all sampling fractions, DivQuant provides the most accurate entropy estimates. In contrast, the observed sample entropy, all asymptotic entropy estimates, and the extrapolating estimator iNext underestimate the true entropy for small sampling fractions and start to overestimate the entropy for larger sampling fractions. On the microbiome data, the entropy confidence intervals of DivQuant are accurate, i.e., DivQuant’s lower bound is smaller than the true entropy and DivQuant’s upper bound is larger.

**Figure 13.**
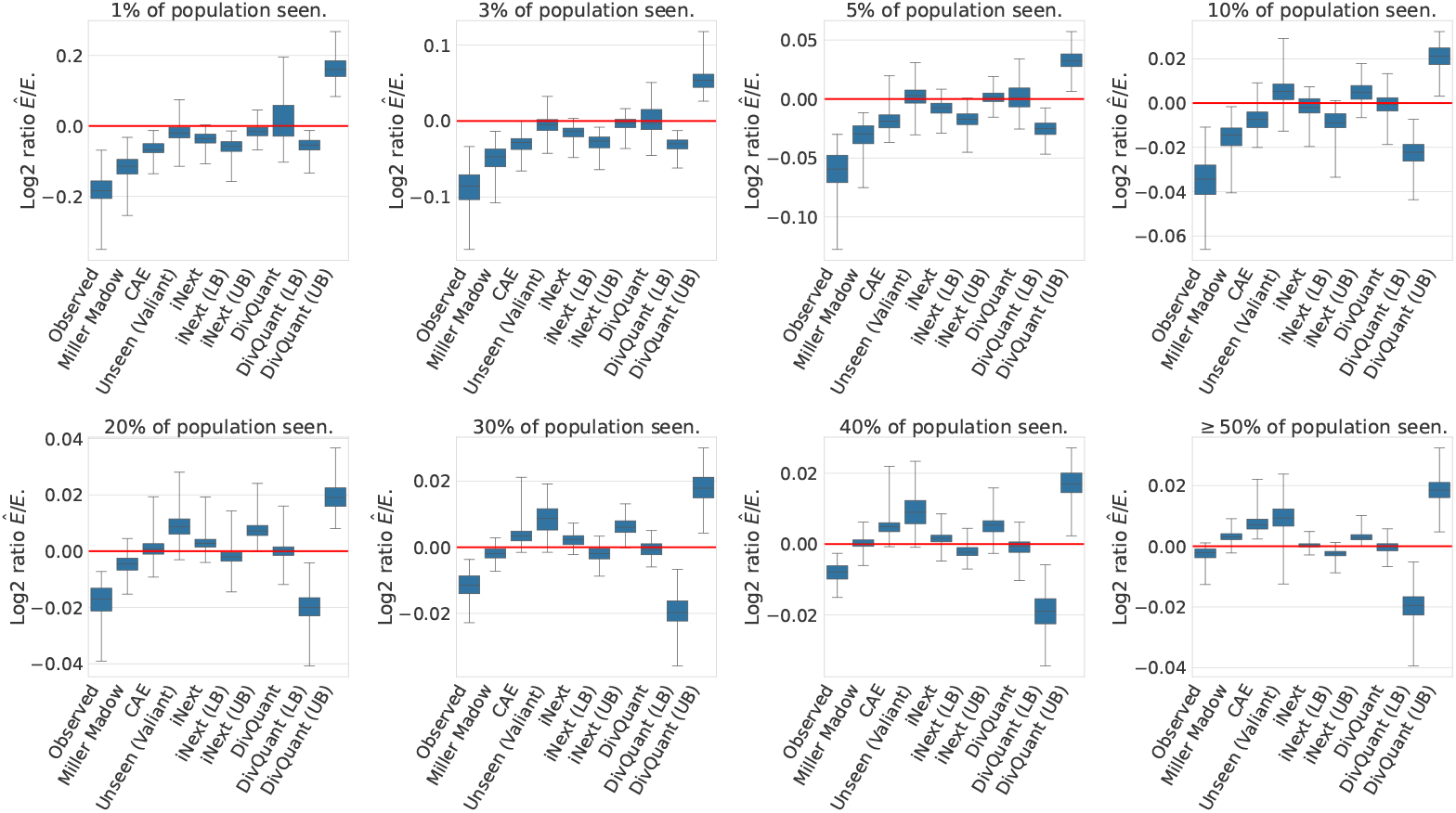
Log_2_ ratio between estimated entropy and true “population” entropy on the *Tara Oceans* microbial taxonomic profiles for different sampling fractions. A log_2_ ratio of zero corresponds to a perfect correspondence, while log_2_ ratios *<* 0 underestimate the true entropy and log_2_ ratios *>* 0 overestimate the true entropy.

## References

1 Alan Agresti and Euijung Ryu. Pseudo-score confidence intervals for parameters in discrete statistical models. Biometrika, 97(1):215–222, 2010. doi:10.1093/biomet/asp074.

2 Steve Baker and Robert D. Cousins. Clarification of the use of CHI-square and likelihood functions in fits to histograms. Nuclear Instruments and Methods in Physics Research, 221(2):437–442, 1984. URL: https://www.sciencedirect.com/science/article/pii/0167508784900164, doi:10.1016/0167-5087(84)90016-4.

3 Kathryn Barger and John Bunge. Objective Bayesian estimation for the number of species. Bayesian Analysis, 5(4):765–785, 2010. doi:10.1214/10-BA527.

4 Joseph Berkson. Minimum Chi-Square, not Maximum Likelihood! The Annals of Statistics, 8(3):457–487, 1980. doi:10.1214/aos/1176345003.

5 Peer Bork, Chris Bowler, Colomban de Vargas, Gabriel Gorsky, Eric Karsenti, and Patrick Wincker. Tara Oceans studies plankton at planetary scale. Science, 348(6237):873–873, 2015. doi:10.1126/science.aac5605.

6 Hendriek C. Boshuizen and Dennis E. te Beest. Pitfalls in the statistical analysis of microbiome amplicon sequencing data. Molecular Ecology Resources, 23(3):539–548, 2023. doi:10.1111/1755-0998.13730.

7 Kenneth P. Burnham and Scott W. Overton. Estimation of the size of a closed population when capture probabilities vary among animals. Biometrika, 65(3):625–633, 1978. doi: 10.1093/biomet/65.3.625.

8 Anne Chao. Nonparametric Estimation of the Number of Classes in a Population. Scandinavian Journal of Statistics, 11(4):265–270, 1984. URL: https://www.jstor.org/stable/4615964.

9 Anne Chao, Nicholas J. Gotelli, T. C. Hsieh, Elizabeth L. Sander, K. H. Ma, Robert K. Colwell, and Aaron M. Ellison. Rarefaction and extrapolation with Hill numbers: a framework for sampling and estimation in species diversity studies. Ecological Monographs, 84(1):45–67, 2014. doi:10.1890/13-0133.1.

10 Anne Chao and Tsung-Jen Shen. Nonparametric estimation of Shannon’s index of diversity when there are unseen species in sample. Environmental and Ecological Statistics, 10(4):429–443, 2003. doi:10.1023/A:1026096204727.

11 Chun-Huo Chiu. A more reliable species richness estimator based on the Gamma–Poisson model. PeerJ, 11:e14540, 2023. doi:10.7717/peerj.14540.

12 Chun-Huo Chiu and Anne Chao. Estimating and comparing microbial diversity in the presence of sequencing errors. PeerJ, 4:e1634, 2016. doi:10.7717/peerj.1634.

13 Robert K. Colwell, Anne Chao, Nicholas J. Gotelli, Shang-Yi Lin, Chang Xuan Mao, Robin L. Chazdon, and John T. Longino. Models and estimators linking individual-based and sample-based rarefaction, extrapolation and comparison of assemblages. Journal of Plant Ecology, 5(1):3–21, 2012. doi:10.1093/jpe/rtr044.

14 Timothy Daley and Andrew D. Smith. Predicting the molecular complexity of sequencing libraries. Nature Methods, 10(4):325–327, April 2013. doi:10.1038/nmeth.2375.

15 Ian A. Dickie. Insidious effects of sequencing errors on perceived diversity in molecular surveys. New Phytologist, 188(4):916–918, 2010. doi:10.1111/j.1469-8137.2010.03473.x.

16 Ronald A. Fisher, Alexander S. Corbet, and Carrington B. Williams. The relation between the number of species and the number of individuals in a random sample of an animal population. Journal of Animal Ecology, 12(1):42–58, 1943. doi:10.2307/1411.

17 Nikolay Gagunashvili. Chi-square tests for comparing weighted histograms. Nuclear Instruments and Methods in Physics Research, 614(2):287–296, 2010. URL: https://www.sciencedirect.com/science/article/pii/S0168900209023547, doi:10.1016/j.nima.2009.12.037.

18 Kate L. Gibson, Yu-Chang Wu, Yvonne Barnett, Orla Duggan, Robert Vaughan, Elli Kondeatis, Bengt-Olof Nilsson, Anders Wikby, David Kipling, and Deborah K. Dunn-Walters. B-cell diversity decreases in old age and is correlated with poor health status. Aging Cell, 8(1):18–25, 2009. doi:10.1111/j.1474-9726.2008.00443.x.

19 Irving J. Good and George H. Toulmin. The number of new species, and the increase in population coverage, when a sample is increased. Biometrika, 43(1):45–63, 1956. doi: 10.1093/biomet/43.1-2.45.

20 François Grin and Guillaume Fürst. Measuring linguistic diversity: A multi-level metric. Social Indicators Research, 164(2):601–621, 2022. doi:10.1007/s11205-022-02934-5.

21 Gurobi Optimization, LLC. Gurobi Optimizer Reference Manual, 2026. URL: https://www.gurobi.com.

22 T. Hauschild and Michael Jentschel. Comparison of maximum likelihood estimation and chi-square statistics applied to counting experiments. Nuclear Instruments and Methods in Physics Research Section A: Accelerators, Spectrometers, Detectors and Associated Equipment, 457(1):384–401, 2001. doi:10.1016/S0168-9002(00)00756-7.

23 Yili Hong. On computing the distribution function for the poisson binomial distribution. Computational Statistics & Data Analysis, 59:41–51, 2013. URL: https://www.sciencedirec t.com/science/article/pii/S0167947312003568, doi:10.1016/j.csda.2012.10.006.

24 T. C. Hsieh, K. H. Ma, and Anne Chao. iNEXT: an R package for rarefaction and extrapolation of species diversity (Hill numbers). Methods in Ecology and Evolution, 7(12):1451–1456, 2016. doi:10.1111/2041-210X.12613.

25 T.C. Hsieh and Anne Chao. Rarefaction and extrapolation: Making fair comparison of abundance-sensitive phylogenetic diversity among multiple assemblages. Systematic Biology, 66(1):100–111, 2017. doi:10.1093/sysbio/syw073.

26 Xiangpan Ji, Wenqiang Gu, Xin Qian, Hanyu Wei, and Chao Zhang. Combined Ney-man–Pearson chi-square: An improved approximation to the Poisson-likelihood chi-square. Nuclear Instruments and Methods in Physics Research Section A: Accelerators, Spectrometers, Detectors and Associated Equipment, 961:163677, 2020. doi:10.1016/j.nima.2020.163677.

27 Krisana Lanumteang and Dankmar Böhning. An extension of Chao’s estimator of population size based on the first three capture frequency counts. Computational Statistics & Data Analysis, 55(7):2302–2311, 2011. doi:10.1016/j.csda.2011.01.017.

28 Daniel J. Laydon, Charles R. M. Bangham, and Becca Asquith. Estimating T-cell repertoire diversity: limitations of classical estimators and a new approach. Philosophical Transactions of the Royal Society B: Biological Sciences, 370(1675):20140291, 2015. doi:10.1098/rstb.2014.0291.

29 Lucien Le Cam. An approximation theorem for the poisson binomial distribution. Pacific Journal of Mathematics, 10(4):1181–1197, 1960. URL: https://msp.org/pjm/1960/10-4/p11.xhtml.

30 George Miller. Note on the bias of information estimates. Information theory in psychology : Problems and methods, 1955.

31 Felix Mölder, Kim P. Jablonski, Brice Letcher, Michael B. Hall, Christopher H. Tomkins-Tinch, Vanessa Sochat, Jan Forster, Soohyun Lee, Sven O. Twardziok, Alexander Kanitz, Andreas Wilm, Manuel Holtgrewe, Sven Rahmann, Sven Nahnsen, and Johannes Köster. Sustainable data analysis with Snakemake. F1000Research, 2021. doi:10.12688/f1000research.29032.1.

32 Jerzy Neyman. Contribution to the theory of the χ2 test. Proceedings of the Berkeley Symposium on Mathematical Statistics and Probability, pages 239–273, 1949. URL: https://projecteuclid.org/ebook/Download?urlid=bsmsp/1166219208&isFullBook=false.

33 Liam Paninski. Estimation of Entropy and Mutual Information. Neural Computation, 15(6):1191–1253, 2003. doi:10.1162/089976603321780272.

34 Nicla Porciello, Ornella Franzese, Lorenzo D’Ambrosio, Belinda Palermo, and Paola Nisticò. T-cell repertoire diversity: friend or foe for protective antitumor response? Journal of Experimental & Clinical Cancer Research, 41(1):356, 2022. doi:10.1186/s13046-022-02566-0.

35 Patrick D. Schloss. Rarefaction is currently the best approach to control for uneven sequencing effort in amplicon sequence analyses. mSphere, 9(2):e00354–23, 2024. doi:10.1128/msphere.00354-23.

36 Johanna E. Schmitz and Sven Rahmann. A comprehensive review and evaluation of species richness estimation. Briefings in Bioinformatics, 26(2):bbaf158, 2025. doi:10.1093/bib/bbaf158.

37 Christopher Schröder and Sven Rahmann. Efficient duplicate rate estimation from subsamples of sequencing libraries. PeerJ PrePrints, 3:e1298v2, September 2015. doi:10.7287/peerj.preprints.1298v2.

38 Claude E. Shannon. A mathematical theory of communication. The Bell System Technical Journal, 27(3):379–423, 1948. doi:10.1002/j.1538-7305.1948.tb01338.x.

39 Gregory Valiant and Paul Valiant. Estimating the Unseen: Improved Estimators for Entropy and Other Properties. Journal of the ACM, 64(6):37:1–37:41, October 2017. doi:10.1145/3125643.

40 Cameron Wagg, Klaus Schlaeppi, Samiran Banerjee, Eiko E. Kuramae, and Marcel G. A. van der Heijden. Fungal-bacterial diversity and microbiome complexity predict ecosystem functioning. Nature Communications, 10(1):4841, 2019. doi:10.1038/s41467-019-12798-y.

41 Pengyang Wang, Donghao Wu, Yongjia Wang, Zhiqiang Shen, Buhang Li, Zufei Shu, and Chengjin Chu. Tree species richness suppresses red imported fire ant invasion in a subtropical plantation forest. Journal of Applied Ecology, 61(11):2751–2761, 2024. doi:10.1111/1365-2664.14786.

42 Amy Willis and John Bunge. Estimating diversity via frequency ratios. Biometrics, 71(4):1042–1049, 2015. doi:10.1111/biom.12332.

43 Amy D. Willis. Rarefaction, Alpha Diversity, and Statistics. Frontiers in Microbiology, 10, 2019. URL: https://www.frontiersin.org/articles/10.3389/fmicb.2019.02407.

44 Jens Zentgraf and Sven Rahmann. Fast Gapped k-mer Counting with Subdivided Multi-Way Bucketed Cuckoo Hash Tables. In Christina Boucher and Sven Rahmann, editors, 22nd International Workshop on Algorithms in Bioinformatics (WABI 2022), volume 242 of Leibniz International Proceedings in Informatics (LIPIcs), pages 12:1–12:20. Schloss Dagstuhl – Leibniz-Zentrum für Informatik, 2022. doi:10.4230/LIPIcs.WABI.2022.12.

